# Dissecting organoid-bacteria interaction highlights decreased contractile force as a key factor for heart infection

**DOI:** 10.1101/2025.04.08.647814

**Authors:** Anheng Wang, Jiaxian Wang, Zhe Zhang, Chuan Yang, Chunhao Deng, Guokai Chen, Chengwu Li, Qian Wang, Lei Dong, Chunming Wang

## Abstract

Bacterial endocarditis is a fatal cardiovascular disease exacerbated by weakened heart contraction, yet the direct impact of cardiac contractility on bacterial adhesion remains elusive. Here, we present a novel quantitative physics model integrating finite element analysis and live-cell imaging to uncover their strong correlation. Using this model, we quantified the real-time force magnitude generated by organoid-type cardiac microtissue derived from healthy donors and dilated cardiomyopathy patients – mimicking normal and suppressed heart contractility, respectively – to the approaching bacteria in a real fluidic system. The data revealed that weakened cardiac contractility facilitated bacterial invasion of the myocardium. Verifying this finding in a mouse transverse aortic constriction model demonstrated that increasing heart contraction efficiently mitigated bacterial invasion, with a 25% increase in heart contractility reducing endocarditis risk by 80%. Our findings demonstrate that patient-derived cardiac organoids provide a physiologically relevant platform for studying bacterial infections in vitro, offering high clinical fidelity. This platform establishes a valuable tool for drug screening and the development of novel therapeutic strategies.

## Introduction

Infective endocarditis (IE) poses significant challenges to global health, affecting 1.5 million patients annually with an over 40% mortality rate[1]. A substantial cause of IE is a bacterial infection, but how the heart becomes prone to the infection remains a fundamental question[1a, 2]. Decades of clinical evidence reveal that patients with a history of valvular heart disease, congenital heart disease, previous infective endocarditis, or cardiomyopathy have a significantly greater chance of IE[3]. All these diseases have the comorbidity of heart failure, as characterized by decreased myocardial contractility[4]. Bacteria must adhere to tissue surfaces for infection[5], and its adhesion is dictated by the status of the surface[6] (e.g. dynamic[7] or static[8]) and the physical environment (e.g. fluidic stress[9]). However, it remains unknown how the heart tissue – under dynamic, repeated, intrinsically driven contractions – influences bacterial adhesion. We hypothesized that decreased myocardial contractility facilitates bacterial infection – which is typically resisted by the robust contractile force in a healthy heart.

Verifying this hypothesis requires a quantitative analysis of the dynamic interactions between the contracting myocardium and moving bacteria, which is challenging for two reasons. The first is the lack of an *in vitro* three-dimensional (3D) cardiac model recapitulating the weakened contractility, as it is unfeasible to adjust the contractility in patients and perform real-time sampling of bacteria-infected hearts *in vivo*. Although organoids have emerged as a powerful 3D tool to mimic various human tissue[10] and organs[11],[12], – including cardiac organoids (COs) of varying extents of maturity[13], they are made of healthy myocardial or progenitor cells and do not exhibit pathologically weakened contractility[11]. The second challenge lies in quantifying bacteria-organoid interaction where COs keep generating intrinsic contractions[14]. Previous analyses have examined regional variations in the contractility of cardiomyocytes[15]. However, it remains challenging to ascertain the physical environment and contractile forces of individual cardiac organoids solely through image recognition techniques. To accurately reconstruct the fluid field of COs and quantify the magnitude of forces exerted by COs on bacteria, a more realistic and sophisticated system is required. Such a system would enable the transformation of motion data into meaningful force measurements, providing deeper insights into the mechanical interactions between COs and their microenvironment.

To overcome these challenges, we aimed to develop an engineering model that integrates human clinical samples with new quantitative tools to dissect how myocardial contractility influences bacterial infection of the myocardium. We first constructed two types of clinically relevant COs through the direct assembly of three cell components into cavity-containing microtissues. Each microtissue comprises human induced pluripotent stem cells (hiPSCs) derived from either patients with dilated cardiomyopathy[16] (DCM) or healthy donors, assembled with both stromal and endothelial cells. These COs were co-cultured with two representative pathogens commonly inducing IE[17]. Distinct from the conventional high-risk IE patients, DCM patients do not exhibit structural cardiac abnormalities similar to congenital heart disease[18], but frequently present with heart failure[19]. These properties make DCM well-suited for investigating whether pure heart failure alone can influence the outcomes of bacterial infections. Both organoids exhibited the dynamic diastole-systole cycle, and the former recapitulated the weakened motility – a key feature of DCM observed clinically. To analyze the interactions in this dynamic setting between planktonic bacteria and beating COs[20], we created a series of quantitative models by applying finite element modelling (FEM)[21] for dynamic live-cell image analysis, which highlighted mechanical features of COs in the fluidic system[22]. Surprisingly, the data revealed a strong inverse correlation between cardiac contractility and bacterial adherence to myocardial tissue. As the contractile force of the cardiomyocyte increased, the number of bacteria adhering to the heart muscle significantly decreased (**Fig. 1**). This effect was consistent across multiple bacterial strains tested. This groundbreaking discovery opens up new avenues for the development of innovative therapeutic strategies to combat cardiac infections. This organoid-based model represents a reliable platform for quantifying the biomechanical forces in bacteria-organoid interactions, thereby opening new avenues for drug discovery and therapeutic innovations.

**Figure 1.**
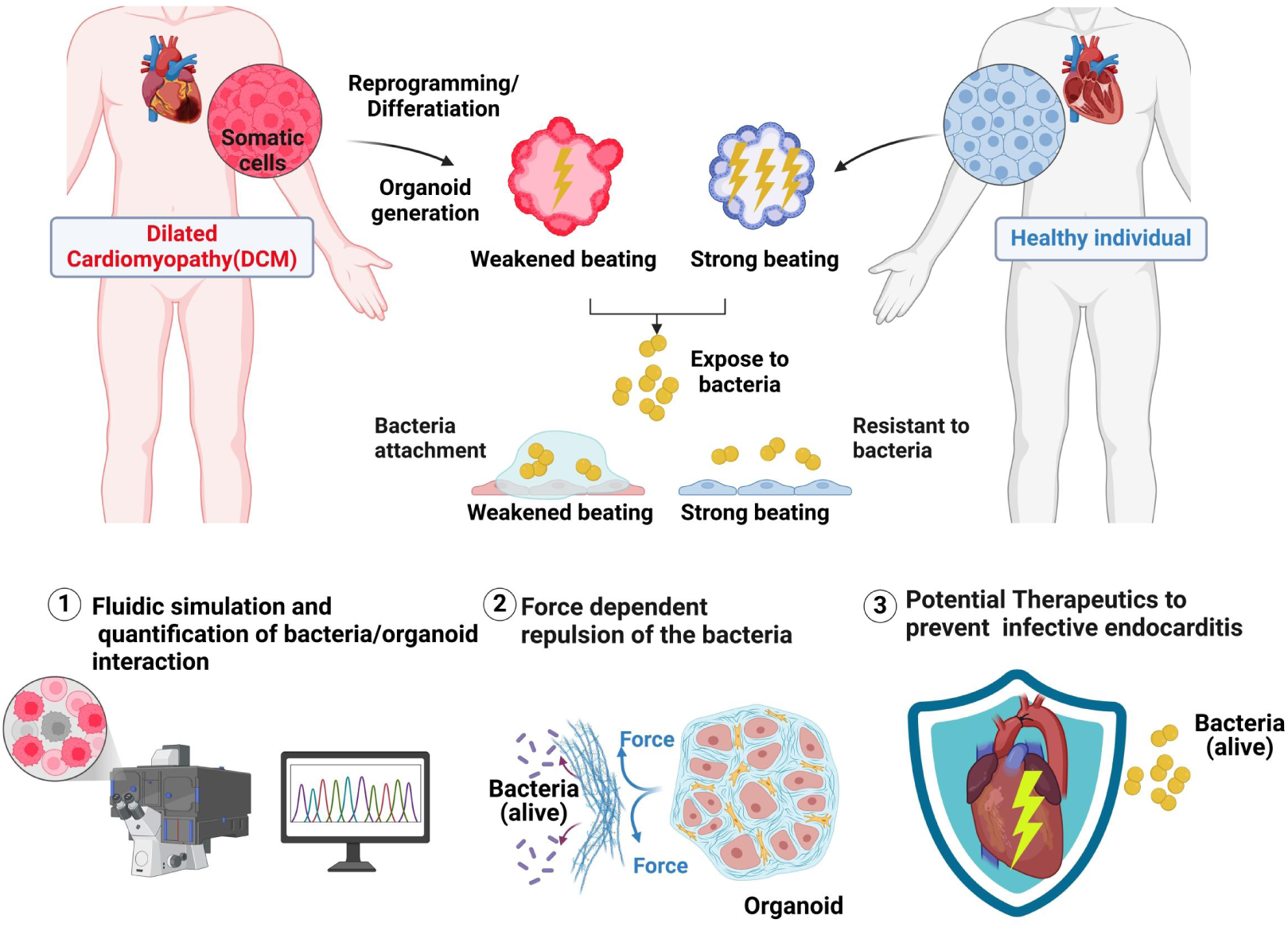
Schematic of experiments with patient-derived cardiac organoids for bacterial infection study.

## Results

### Construction of COs exhibiting pathologically decreased contractility

To construct 3D organoids recapitulating regular and weakened heart contractions, we first generated two clinically relevant CMs in 2D monolayers from hiPSCs derived from DCM patients and healthy individuals – namely DCM-CMs and H-CMs, respectively, following an established protocol[23] (**Fig. 1**). DCM-CMs and H-CMs exhibited different morphologies and Troponin localization (**Fig 2A**; staining for troponin on day 33) and had high proportions of cardiomyocytes exceeding 85% (DCM, **Fig. S1A)** and 97% (CM. **Fig. S1B**), respectively. Similar to its corresponding clinical symptoms, DCM showed significantly weaker contractility compared with CM, as evidenced by patch clamp tests (**Fig. 2B)**, although their beating rates were similar (**Fig. 2C**). The DCM cells were derived from a 47-years-old Asian male with an LMNA gene mutation. The LMNA gene primarily encodes lamin A and C, which are nuclear lamina proteins; its mutation can lead to dilated DCM, mainly by impairing cardiac contractile function and causing cardiac conduction block[24]. Our data indicated that the DCM and CM had distinct transcriptional profiles (**Fig. S1C**; bulk RNA sequencing [RNA-seq] on day 22). Compared with H-CMs, DCM-CMs expressed significantly lower levels of MYH7 (for cardiac cell contraction) and FLNC (for maintaining sarcomere integrity), among other contractility-related genes (e.g. ACTN2, HCN4, TNNI1), and a higher level of PLN (associated with arrhythmogenic in DCM patients[25]).

**Figure 2.**
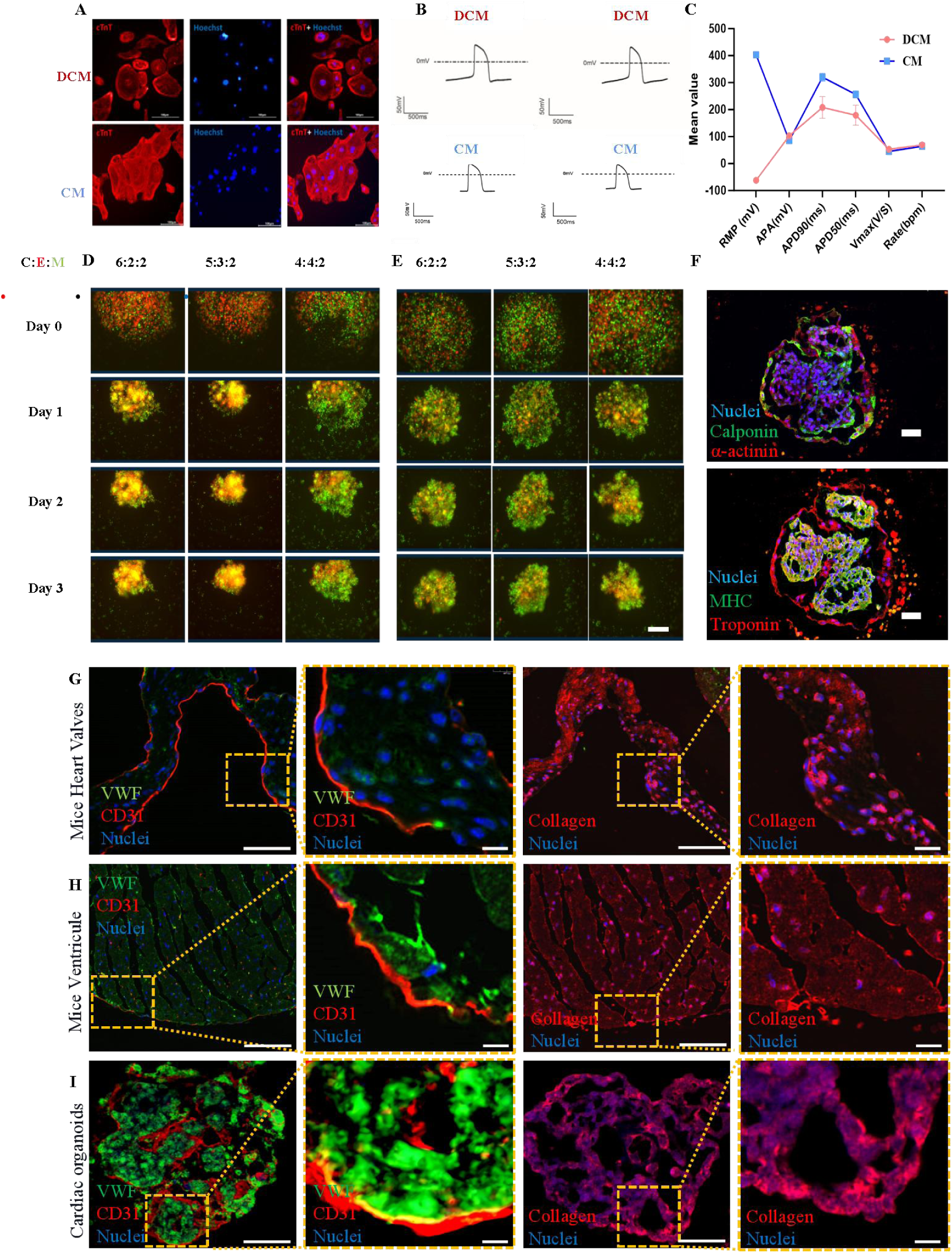
Preparation of human cardiac organoids (COs) of healthy and dilated cardiomyopathy (DCM) characteristics. (A) Immunohistochemical staining for cardiomyocyte biomarker (Troponin). Scale bars, 100 µm. (B) Electrophysiological data of DCM and CM measured using patch clamp. (C) Membrane patch-clamp measurements of the potential and heartbeat frequency data in DCM and CM. (D) Fluorescence microscope observation of the formation of HCOs with different cell ratios. The ECs were red (Celltracker-far red), and MSC was green (CFSE) in the Fluorescence Observation. The bar is 50µm. (E) Fluorescence microscope observation of the formation of DCM-COs with different cell ratios. The ECs were red (Celltracker-far red), and MSC was green (CFSE) in the Fluorescence Observation. The bar is 50µm. (F) Confocal Observation of Calponin / α-actinin / MHC / Troponin localisation within the HCOs. The Calponin / MHC were labelled green (Alexaflour 488), and Troponin / α-actinin were labelled red (Alexaflour 647). The bar is 50 µm. (G) Immunohistochemical staining for classic endocarditis biomarker (vWF, CD31 and Collagen Type 1) on Mice heart valves section. The vWF waslabelled green (Alexaflour 488), CD31 Collagen Type 1 and was labelled red (Alexaflour 647). Scale bars, 10µm. (H) Immunohistochemical staining for classic endocarditis biomarker (vWF, CD31 and Collagen Type 1) on Mice heart ventricle section. The vWF waslabelled green (Alexaflour 488), CD31 Collagen Type 1 and was labelled red (Alexaflour 647). Scale bars, 10µm. (I) Immunohistochemical staining for classic endocarditis biomarker (vWF, CD31 and Collagen Type 1) on Cardiac organoids section. The vWF waslabelled green (Alexaflour 488), CD31 Collagen Type 1 and was labelled red (Alexaflour 647). Scale bars, 10µm.

We performed cell movement analysis based on over 50 plots using TrackMate (Jean-Yves Tinevez and Jaqaman et al. [26]). Both CMs started to develop contractility on day eight at ∼ 50-70 beats per minute (bpm), close to the adult human heart rate of 60-100 bpm (**Fig. S1D**). H-CMs contracted with a constant intensity similar to a healthy heart (**Video S1**), while DCM-CMs displayed an irregular frequency and amplitude of exercise with reduced intensity (**Video S2**).

Then, we constructed 3D microtissues towards assembly into cavity-containing COs, based on these 2D CMs and DCMs. Physiologically, the endocardium comprises CMs, endothelial cells (ECs), and stromal cells. Co-culturing CMs with both human umbilical vein endothelial cells (HUVECs, representing ECs) and human mesenchymal stem cells (MSCs, representing stromal cells) formed larger, more compact spheroid microtissues than culturing CMs alone or CMs with ECs (**Fig. S1E**). We tested three different proportions for the tri-cellular composition of CM and DCM, respectively (**Fig. 2D** and **E**) and found the optimal one for constructing the 3D microtissue was 6:2:2 (CM/DCM: EC: MSC). Under this condition, the spheroidal microtissue aggregated more rapidly and exhibited stronger motility than under others, similar to their actual proportion in the human fetal heart. The addition of MSCs and ECs to CMs increased the expression of cell motility-and structure-related genes (e.g. ACTG1, COL4A2, COL5A3, TIMP12; **Fig. S1F**). However, in MSC/ECs/CMs co-cultured organoids, MLF1 and SCP2 were down-regulated compared to individual CMs culture, which accounts for hematopoietic cell development lipid metabolism, respectively. In this ratio, the ECs formed apparent networks spreading throughout the microtissue after five days of culture (**Fig. 2F, S1G)**, with a live recording of organoid formation in this cellular proportion from Day 1 through Day 3 (Red, ECs; Green, CM; **Video S3**).

Next, we examined these microtissues for their characteristics as cavity-containing COs. Indeed, they contained clear cavity structures – as evidenced by morphological observation upon MHC and troponin staining for myosin (**Fig. 2F**), which are highly comparable with other studies on the construction and definition of COs[13a, 27]. Further examination of the different layers from top to bottom of these COs demonstrated the presence of substantial cavities in the core of these organoids (**Fig. S1G**). Thus, we adopted this condition to prepare DCM-COs and HCOs in subsequent experiments.

Finally, we determined the suitability of these COs as a model for endocardial bacterial infection, by staining for three key targets involved in *Staphylococcus aureus* (*S. A*) infection – vWF, CD31, type 1 collagen[24, 24, 28] – in mouse heart sections and our engineered HCOs in parallel. We noticed a consistent distribution of all three markers between them (mouse heart valves, **Fig. 2G**; mouse ventricular endocardium, **Fig. 2H**; HCOs, **Fig. 2I**). Specifically, CD31 and vWF were primarily distributed on the surface and in a deeper site, respectively, of the tissue, while Type I collagen spread widely.

### Characterization of the contractility of the two clinically-derived COs

We examined the contractility of the two CO types by dividing an organoid into multiple regions at 15-μm intervals and using TrackMate^®^ to track the precise motion of individual regions (**Fig. 3A)**. We observed a higher amplitude of contraction in HCOs (**Video S4**) than in DCM-COs (**Video S5**), specifically including a higher mean speed (**Fig. 3B, S2A)**. Meanwhile, DCM-COs exhibited a clear central motion with weaker motility on the edge, while HCOs’ contraction was relatively uniform (**Fig. 3C, S2B**). Similarly, the straightness of DCM-COs’ motion was lower, as its movement was less forceful or regular (**Fig. S2C**). These findings agreed with the outcome observed in 2D CMs.

**Figure 3.**
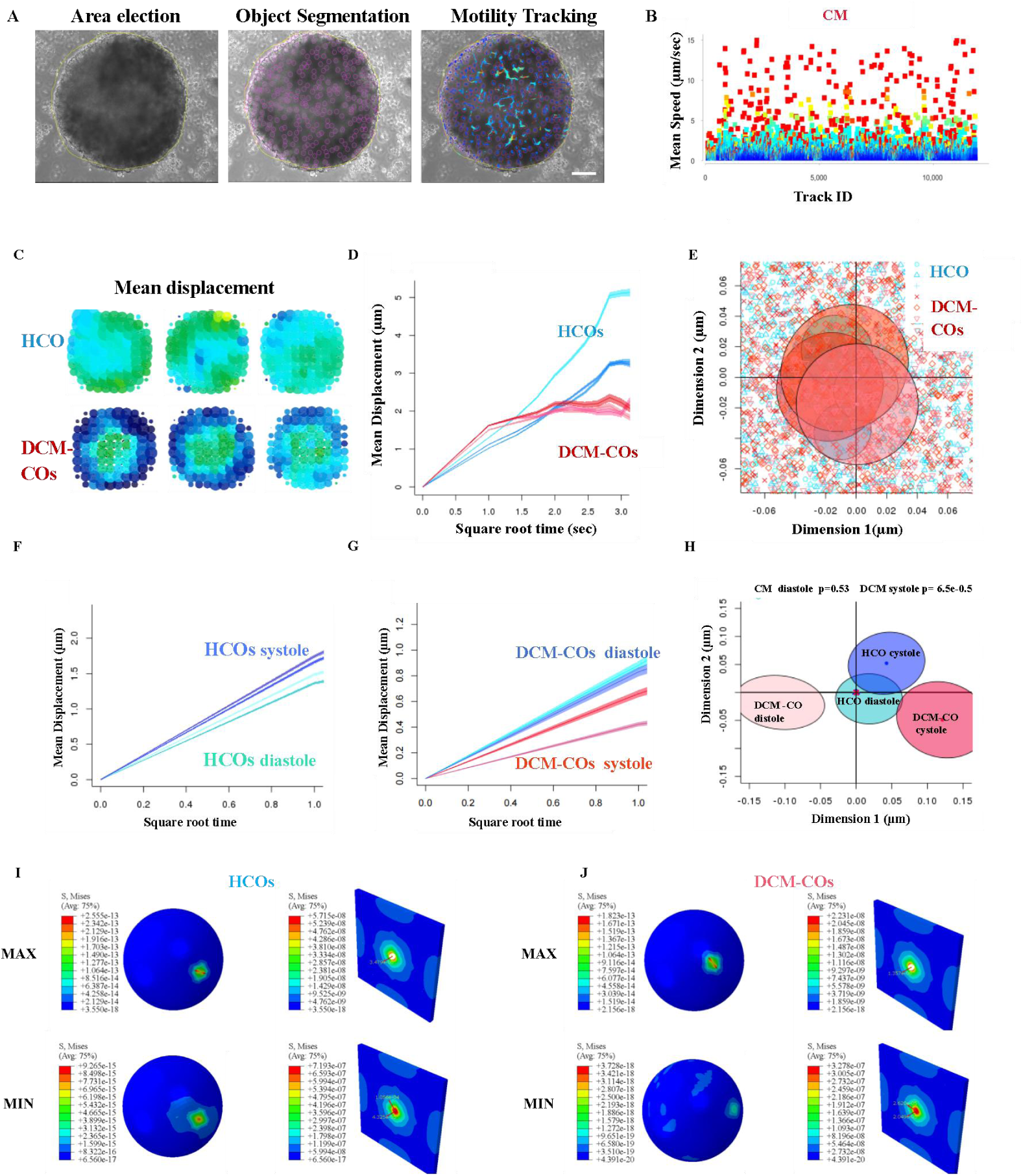
Dissection of COs-bacteria interactions in the contraction-induced fluidic system. (A) Tracking process of the COs movement. The first step was to check the tracking area, the second was to confirm the partition size, and the third was to select the appropriate threshold for tracking. (B) Mean speed for different tracked IDs obtained from HCOs. Each CO was divided into 100-200 15-μm circles. (C) Mean displacement heatmap of the HCOs and DCM-COs. Each CO was divided into 100-200 15-μm circles. Tested on two group of 6 independent COs in total (n=6). (D) The total displacement produced by the HCOs and DCM-COs in 3 seconds. (E) Hotelling’s Test examines the correlation between HCOs and DCM-COs moving in 3 seconds. (F) Displacement of HCOs within a single systolic-diastolic cycle for contraction and circulation, respectively. Tested on two groups of 6 independent HCOs in total (n=6). (G) Displacement of DCM-COs within a single systolic-diastolic cycle for contraction and circulation, respectively. Tested on two groups of 6 independent DCM-COs in total (n=6). (H) Two-dimensional hotelling’s test analysis of motion correlations between HCOs and DCM-COs for systolic and diastolic displacement processes. (I) Finite element analysis of the forces exerted by HCOs on a 1-µm^2^ water field. The analysed velocities of the acting forces were taken from the maximum and minimum velocities of the HCOs obtained from the tracking procedure. (J) Finite element analysis of the forces exerted by DCM-CO on a 1-µm^2^ water field. The analysed velocities of the acting forces were taken from the maximum and minimum velocities of the DCM-COs obtained from the tracking procedure.

We recorded that the mean displacement of HCOs through two systole-diastole cycles was 2.5 times greater than that of DCM-COs (**Fig. 3D**). For each completed motion, we performed Hotelling’s test for three different COs. The within-group range of DCM-COs was more comprehensive, indicating poorer internal consistency. At the same time, the overall motion patterns of HCOs and DCM-COs were similar (**Fig. 3E**), with no significant difference (*P*>0.05). However, if the complete motion was divided into systole and diastole processes, HCOs, like a normal heart, exhibited slightly greater contraction than relaxation (**Fig. 3F**). In contrast, DCM-COs showed the opposite pattern, with higher relaxation than contraction (**Fig. 3G**). The 2D spatial motion tests also indicated that the two segments of motion in DCM-COs were utterly uncorrelated, while HCOs exhibited a considerable degree of correlation (**Fig. 3H)**, as also recorded by a live-transformed tracking video (**Video S6**; red, HCOs; blue, DCM-COs). DCM-COs reproduced the characteristics of the disease – an imbalance between systolic and diastolic blood pressure. HCOs showed a systolic cycle similar to a normal heart, with their systolic pressure greater than diastolic pressure[26a, 29]. Overall, HCOs and DCM-COs replicated the patterns of disease movement in the body, with DCM-COs exhibiting weaker motion characteristics and contraction-release dysregulation.

### FEM analysis of CO-bacteria interactions in the contraction-induced fluidic system

The contractile force of DCM exceeds that of relaxation, which may result in bacteria being more likely to approach the edges of COs, potentially facilitating bacterial adhesion. To verify the validity of this hypothesis, we initially constructed a simulated fluidic system to model the behavior of particles near DCM-CO and CM-CO interfaces.

i) setting appropriate thresholds to acquire parameters (e.g. velocity of motion and average displacement),
ii) transforming the coupled data of the Fourier series into a sinusoidal motion waveform and inputting the sinusoidal waveform into the fluid system to construct a sphere-like motion and
iii) using the FEM model to simulate the movement of particles near the sphere (Representing COs) to make an initial inference on the behaviour of bacteria in the vicinity.

The region-specific motility (velocity) generated above enabled us to evaluate the force in the fluid-solid system that can affect bacterial adhesion. We employed FEM to analyse how the contracted COs could influence the surrounding fluidic environment and bacterial adhesion in a real, dynamic physical system. The results showed that “points” (referring to the velocity of individual zones tracked by TrackMate^®^) appearing in the high-speed segment (red) were more in HCOs (**Fig. 3B**) than in DCM-COs (**Fig. S2A**); over 70% of points fell within the low-speed motion range in the latter. We extracted the data of COs contraction and converted them reversely through Fourier-transformed into motion curves in the Ansys fluent system[30].

As illustrated in **Fig. S2D** and **E**, a white circle in the center represents a fit CO derived from TrackMate-generated data of HCOs or DCM-COs. The water around HCOs exhibited a higher flow velocity than that around DCM-COs because of the more muscular contraction of the former. In the FEM system, we “placed” 1-µm-particles of “bacteria” at different distances from the organoid center and employed the ANSYS Fluent software to predict the trajectory of a “bacteria cell” by integrating the force balance on the particle calculated by a Lagrangian reference frame. This force balance equates the particle inertia with the forces applied to the particle, as written below:

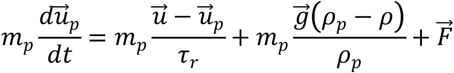

Where mp is the particle mass, is the fluid phase velocity, up is the particle velocity, and p is the liquid density. P is the density of the particle, F is an additional force, 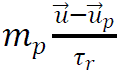 is the drag force, and Tr is the droplet or particle relaxation time calculated by:

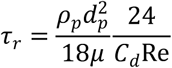

Here, μ is the molecular viscosity of the fluid, dp is the particle diameter, and Re is the relative Reynolds number, which is defined as

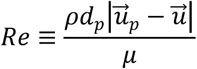

The “bacteria” at different distances received varying forces, with those closer to the organoid boundary subjected to more excellent propulsion. Since the input mass data, density, and other parameters of bacterial particles and cells are consistent, the primary distinguishing factor for particle motion is the motion data of the COs.The particles around HCOs moved in a similar pattern as the diastole-systole cycle; those almost on the organoid boundary were slightly pulled closer at the end of each cycle, while those at a greater distance almost remained in place. Notably, HCOs (**Fig. 3I**) could generate 1.5-time higher maximum force (43.25 µN versus 27.76 µN) and nearly 5-time greater minimum force than did DCM-COs (**Fig. 3J**), which should pose a significant impact on the bacteria. However, by measuring the positions of the “bacteria” and COs at the beginning and end of their motion (**Fig. S2F** and **G**), we observed that the contraction and relaxation of COs generated two forces on the bacteria – pushing and pulling – that eventually failed to push the bacteria away from the organoid boundaries. Different motion intensities of HCOs and DCM-COs led to varying influences on the bacteria. HCOs, with regular systolic and diastolic cycles, possessed higher forces and caused stronger absolute displacements of particles – but paradoxically brought bacteria closer. DCM-COs, resulting from abnormal systolic and diastolic cycles, pushed bacteria further away. To solve this contradiction, verifying which factor (final coordinate or mean displacement) ultimately has a greater influence on bacterial adhesion is necessary.

### Quantitative elucidation of decreased CO contractility on increased bacterial infection in co-culture

We cultured COs with *S. A*, the most common bacteria species inducing IE clinically. This species cannot move via self-propulsion due to their lack of flagella or cilia [31] and exhibit Brownian motion. Based on their weak motility, we speculated that they, similar to the particles in the FEM system, could be strongly influenced by the COs. To validate this assumption, we placed these two bacteria near the contracting COs and tracked them for an extended period of 48 hours.

We immediately tracked the movement of *S. A* within 100 μm to the periphery of HCO (**Fig. S2H**) or DCM-CO (**Fig. S2I**) after inoculation. The bacteria around HCO formed an elongated straight line (**Fig. 4A**), markedly different from the point-like Brownian motion observed in the control group without COs (**Video S7, Fig. 4C**). The weak motility of DCM-COs possessed a much slighter influence on the movement of bacteria than HCOs did (**Fig. 4B, Video S8**). Upon thorough analysis of a single bacterial movement near HCO (**Fig. 4D)** or DCM-CO (**Fig. 4E)**, it was observed that bacteria experienced stretching and pulling due to the contraction and relaxation of COs, with a synchronized, oscillatory motion. Moreover, the experimental results were highly consistent with the simulations. After completing a cycle of contraction and relaxation of COs, the final position of bacteria indeed tended to be closer to COs, whereas the results around DCM-COs were contrary (**Fig. 4D and E**).

**Figure 4.**
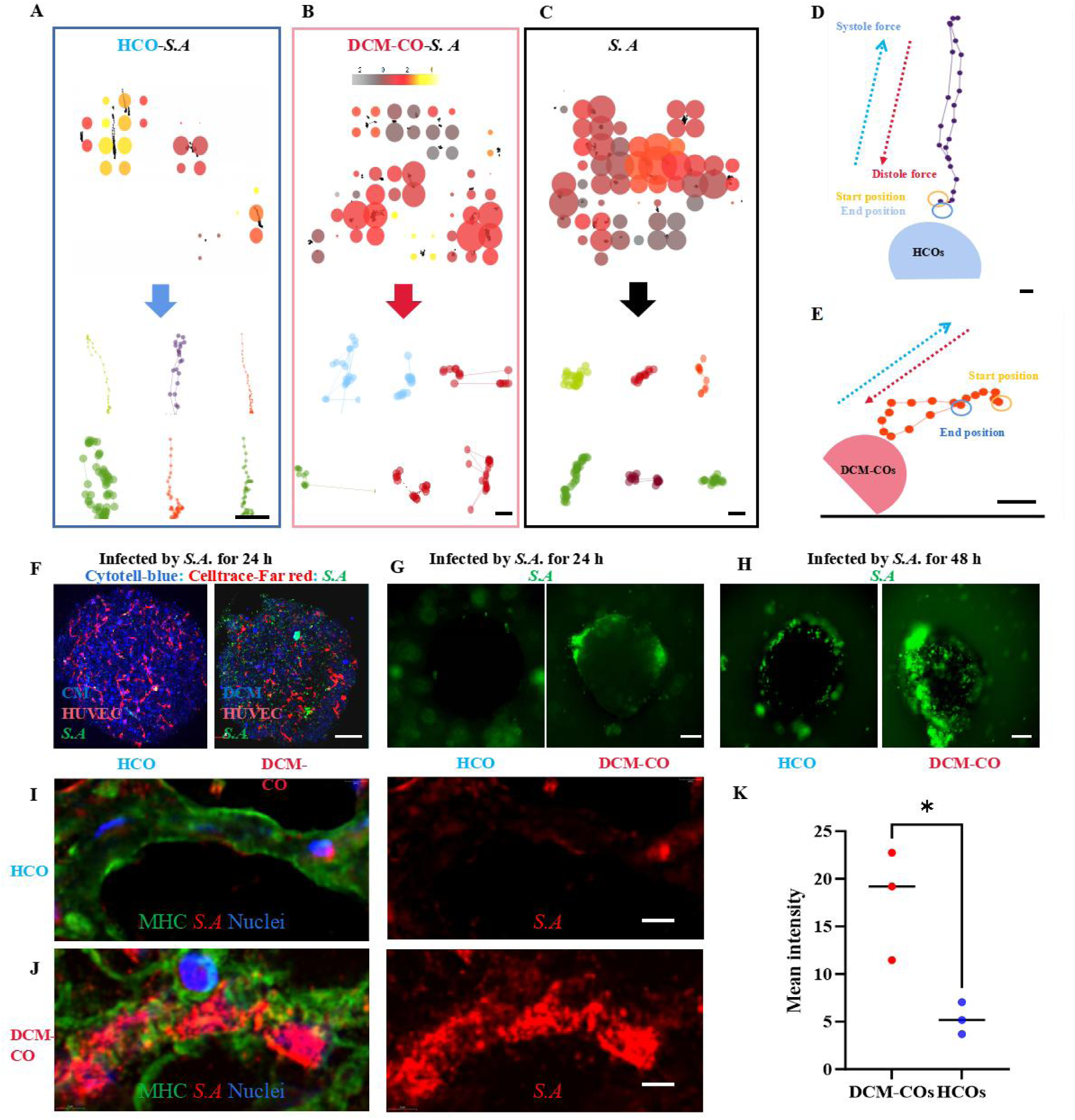
Dynamic analysis of bacterial infection of cardiac organoids. (A) Heatmap of the total displacement and track path of *S. A* around HCOs. The bar is 50 μm. (B) Heatmap of the total displacement and track path of *S. A* around DCM-COs. The bar is 10 μm. (C) Heatmap of *S. A* total displacement and track path of the planktonic *S. A* in the medium. For (A-C), the tracing lasting for two diastolic-systolic cycles was 3 seconds. The bar is 10 μm (D) Analysis of the movement routes of *S. A* in the systole-diastolic cycle near the fringe around HCOs. The bar is 10 μm (E) Analysis of the movement routes of *S. A* in the systole-diastolic cycle near the fringe around DCM-COs. The bar is 10 μm (F) Z-stacked live observation of *S. A* infection after a 24-hour co-culture with HCOs (left) or DCM-COs (right). Red: HUVEC (CellTrace Far-red); blue: CM (Cytotell Blue); Green: *S. A* (CFSE). The bar is 50 μm (G) Fluorescence observation of *S. A* infection after a 24-hour co-culture with HCOs (left) or DCM-COs (right). Green: *S. A* (CFSE). The bar is 50 μm (H) Fluorescence Observation of the infection outcome of *S. A* infection after 48h co-culture with HCOs and DCM-COs, respectively. Green: *S. A* (CFSE). The bar is 50 μm (I) CLSM Observation MHC/DAPI/*S. A* of the infection outcome of *S. A* infection after 24h incubation in the vicinity of HCOs.The bar is 10 μm (J) CLSM Observation MHC / DAPI / *S. A* of the infection outcome of *S. A* infection after 24h incubation in the vicinity of DCM-COs.The bar is 10 μm (K) Quantification of the bacterial attachment on HCOs and DCM-COs after 24h incubation using fluorescence intensity measurement.

After an additional 24 hours, we observed the adhesion of *S. A*. Surprisingly, a magnificent amount of bacteria adhered to DCM-COs after 24 hours of inoculation, whereas very few adhered to HCOs (**Fig. 4F, G**). Following an additional 48 hours of co-incubation, both types of COs were noticeably adhered to by *S. A* (**Fig. 4H**). We co-localized MHC distribution in *S. A* and COs after 24 hours of co-culturing. The *S. A* adhered to them significantly less compared to DCM-COs (**Fig. 4I, J,K**).

These results seemed to contradict our previous simulations and motion analyses, suggesting that the motion of COs does not necessarily make bacteria more susceptible to infection by bringing them closer. Notably, from 24 to 48 hours, the bacteria started adhering to HCOs, accompanied by a markedly decreased motility with these organoids (**Video S9**). Also, bacteria disrupted the overall movement of COs, which changed towards local peristalsis 48 h post-infection (**Video S10**), indicative of the significantly weakened motility of COs. Based on these results, we speculated that COs’ motility could influence the infection outcomes instead of the diastole-systole imbalance, as HCOs exhibited significantly stronger motility than DCM-COs.

To determine whether the motility of COs can deviate from bacterial adhesion, we run trajectory and motion analysis on the bacteria around the COs. First, we analyzed the overall motion trajectories of CO and nearby *S. A*. We found that both HCO (**Fig. 5A**) and DCM-CO (**Fig. 5B**) could change the motion pattern of surrounding *S. A* from Brownian motion to more ordered motion. *S. A* at the edge of HCOs was entrained by the movement of HCOs, resulting in an average displacement even 1.5 times greater than that of HCOs themselves (**Fig. 5A)**, while the displacement near DCM-COs remained the same with that of the HCOs (**Fig. 5B**). Overall, the displacement of *S. A* around HCOs was more than twice that around DCM-COs. T Hotelling’s test also indicated that the *S. A* near CM was utterly unrelated to that of planktonic *S. A*, while *S. A* around DCM-COs remained related (**Fig. 5C**). Notably, although the motion patterns of the two CO types partially overlapped, those of *S. A* around each CO type were distinct (**Fig. 5D**). The reason might be that HCO, representing the healthy tissue, exerted twice as strongly as DCM-COs, reflecting the pathological myocardium, to alter bacterial motility to resist infection.

**Figure 5.**
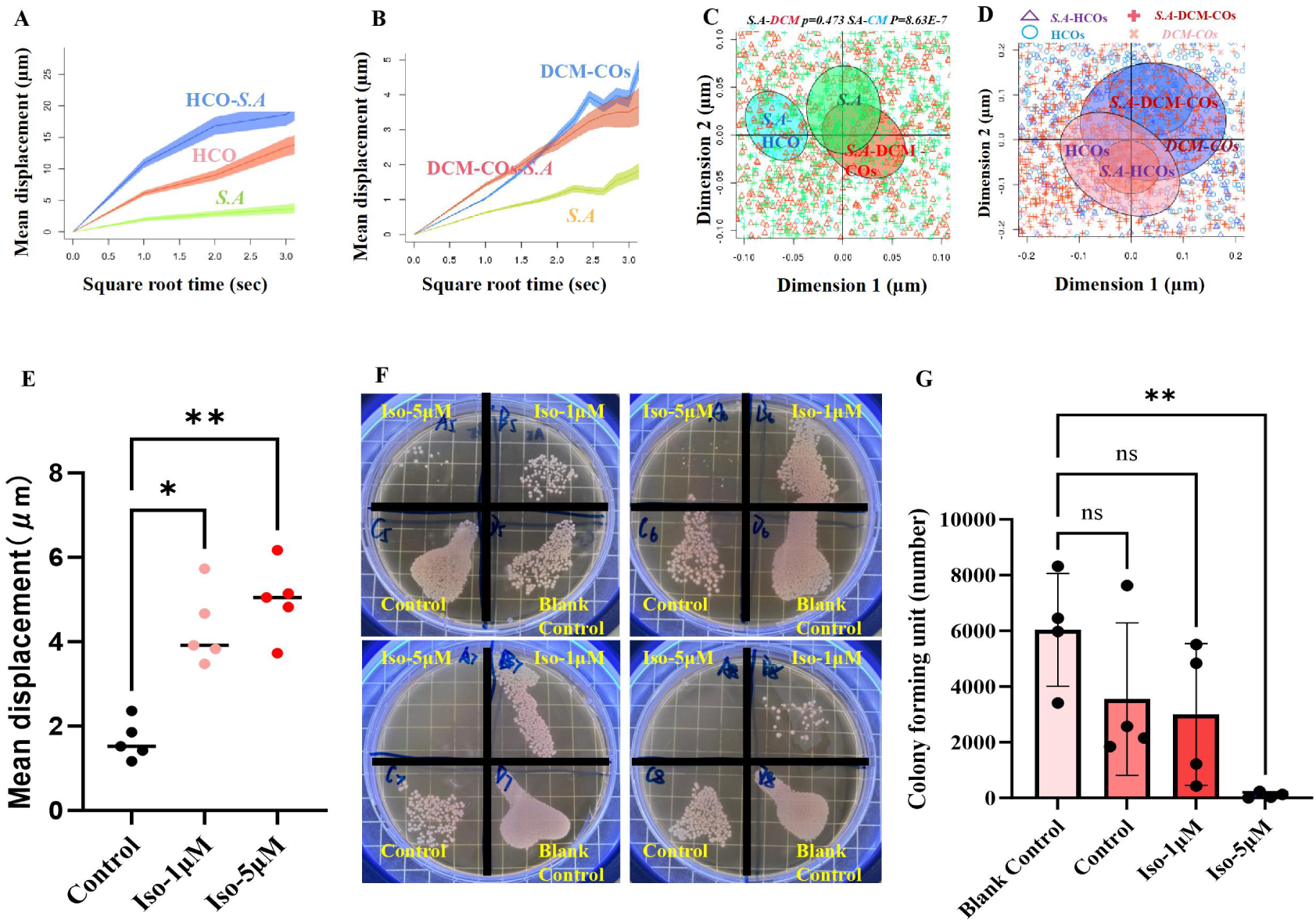
Quantifying the influence of force on bacteria adhesion. (A) Mean displacement of *S. A* near HCOs (HCO-*S. A*) and correlated HCOs compared to planktonic *S. A*. (B) Mean displacement of *S. A*-near DCMCOs (DCMCO-*S. A*) and correlated DCMCOs compared to planktonic *S. A*. (C) Two-dimensional hotelling’s test analysis of motion correlations between *S. A-near* HCOs (HCO-*S. A*), *S. A* near DCM*-*COs (DCMCO-*S. A*) and planktonic *S. A*. (D) Two-dimensional hotelling’s test analysis of motion correlations between *S. A*-near HCOs (*S. A*-CM) and correlated HCOs (HCOs), *S. A* near DCM*-*COs (*S.A*-DCM) and correlated DCM-COs (DCM-COs) (E) Quantitative analysis of HCOs contraction strength (total displacement during a systole-diastolic cycle) after treatment with different concentrations of isoproterenol (1 or 50μM). (F) The infection status of HCOs organoids after 24-hour co-culture with *S. A* (10^5^/mL) under different 3 treatment groups (PBS as control, 1 or 5 μM isoproterenol as drug) compared to the *S. A* in medium (Blank Control) determined by the Colony forming unit (CFU) method. (G) The counting results in the CFU assay obtained from Fig. 5F.

Therefore, the contracting COs appeared to inhibit bacterial adhesion by affecting the latter’s movement. To validate this speculation, we employed two additional bacteria species – *Pseudomonas aeruginosa* (*P. A*) and *Streptococcus anginosus* (*S. S*). The former possesses vigorous motility (up to 10 µm/s) because of its robust flagella, parallel with the other two stationary bacteria. The latter performs like *S. A, exhibiting* Brownian Motion. Indeed, *P. A* exhibited significantly stronger motility (**Fig. S3A, B and C**), with a greater mean displacement (**Fig. S3D**) and speed (**Fig. S3E**) during a 3-second tracking period than *S. A* and *S. S*. Consequently, *P. A* demonstrated a double average velocity (**Fig. S3E**) and four-time larger displacement (**Fig. S3F**) and double average velocity than the other two non-motile bacteria. Meanwhile, the motility of *P. A* was hardly affected by COs due to its strong velocity (**Fig. S3G)**. The two-dimensional Hotelling test further revealed that HCO-*P. A*, DCM-CO-*P. A*, and *P. A* alone had similar patterns of movements (**Fig. S3H**). Interestingly, neither HCOs nor DCM-COs could remain uninfected after 24-hour co-incubation with *P. A* (**Fig. S3I** and **J**), consistent with this bacteria’s motility characteristics observed above. Besides, like *S. A*, *S. S* infected HCOs lightly (**Fig. S3K**) but DCM-COs severely (**Fig. S3L, M**). These results indicated that the motility of the bacteria and COs collectively determined the infection outcomes.

However, was it because of different surface characteristics – instead of motility – between the two COs that made DCM-COs easier for adhesion? To answer this question, we measured the adhesive forces of the bacteria on different COs by coating the bacteria on AFM probes and cantilever (**Fig. S4A**). Bacteria adhered to cantilevers using poly-L-lysine, and the adhesion forces between COs and *S. A* (**Fig. S4B)**, *S. S* (**Fig. S4C)**, or *P. A* (**Fig. S4D)** were measured in the probe mode[32]. The results indicated that *S. A* exhibited greater adhesive forces on HCOs (> 30 nN) than on DCM-COs (**Fig. S4E**); meanwhile, *P. A* showed a similar adhesion to both COs (**Fig. S4F**). These data excluded varying bacterial adhesion to different CO types as the reason for different infection scenarios. Our findings uncovered the correlation between the infection of COs by planktonic bacteria and the motility of COs themselves.

In addition, we tested whether enhancing cardiomyocyte beating could further reduce infection. We accelerated the contractility of HCOs with Isoproterenol (ISO) [33], which, at varying concentrations, significantly enhanced cardiac activity (**Fig. 5E**) and thereby inhibited bacterial adhesion. At 5 uM (**Fig. 5F**), ISO triggered a nearly fourfold increase in beating capacity that resulted in almost no bacterial attachment to the COs (**Fig. 5G**).

### Omics analysis revealing *S. A* disrupts CO contraction to facilitate infection

Based on our previous observations, we identified a correlation between bacterial infection and the motility strength of COs. However, after 48 hours of cultivation, both highly and weakly motile COs failed to escape bacterial adhesion. We speculate that this phenomenon may be attributed to the impairment of COs’ motility following co-cultivation of bacteria.

Based on this hypothesis, we observed that the movement of the HCOs after infection was significantly mitigated after 24h inculcation with *S. A* more than two times (**Fig. 6A**). Our Hottling’s test result also indicated that The motility of infected COs and HCOs becomes significantly divergent (**Fig. S5A**). Alongside these observations, the cellular activity against bacteria also sharply declines to only 4.543N, over nine times lower than the HCOs (41.86N) (**Fig. S5B**).

**Figure 6.**
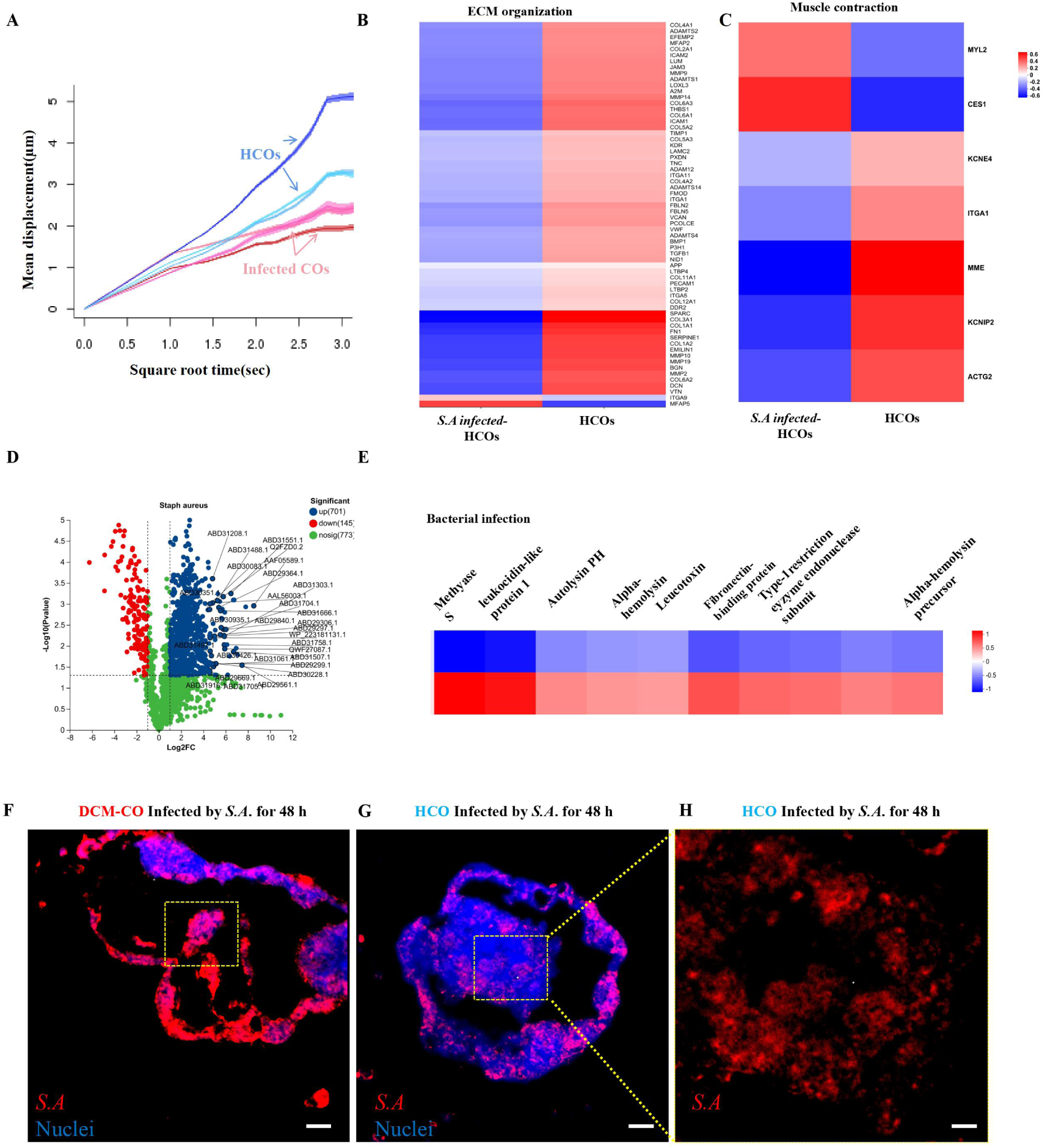
Dissecting the influence of toxins on CO contractility and bacterial infection. (A) Mean displacement of the *S. A* infected HCOs (24-hour inoculation) and the healthy control HCOs, the track length is 3s. n= 6 biologically independent samples (Two groups for HCOs/Infected HCOs each). (B) Heat map of DE genes in the metabolic pathway Reactome ECM organisation Pathway (R-HSA-1474244) in the organoid model showing different trends in the changes in gene expression after *S. A* infection compared to healthy organoid control after 24 hours. The colour scale shows the row z score. Organoid samples, n= three biologically independent samples per group (each sample containing one well of 10-20 organoids); (C) Heat map of DE genes in the metabolic pathway Reactome Muscle Contraction Pathway (R-HSA-397014) in the organoid model showing different trends in the changes in gene expression after *S. A* infection compared to healthy organoid control after 24 hours. The colour scale shows the row z score. Organoid samples, n= three biologically independent samples per group (each sample containing one well of 10-20 organoids); (D) Distribution of protein secretion and fold change; non-DE (yellow) and DE genes (Red for downregulation and blue for up-regulation) of *S. A* isolated from *S. A* HCOs infection model were compared with the control *S. A*. (E) Heat map of DE protein secretion in the metabolic pathway KEGG bacterial infection Pathway (H02076) in the organoid model shows different trends in gene expression changes after *S. A* infection compared to healthy organoid control after 48 hours. The colour scale shows the row z score. Organoid samples, n= three biologically independent samples per group (each sample containing one well of 10-20 organoids); (F) Immunohistochemical staining for *S. A* infection site on H-COs after 24h incubation. The bar is 10 μm. (G) Immunohistochemical staining for *S. A* infection site on HCOs after 24h incubation. The bar is 10 μm. (H) Immunohistochemical staining for *S. A* infection site on HCOs after 24h incubation. The bar is 4 μm.

On the dynamic COs-bacteria interactions, we sought to understand how bacteria influenced CO contraction to increase adhesion. First, we verified that *S. A* inoculation greatly changed the expression of genes related to tissue structure and motility, such as ECM production and degradation and myocardial contraction (**Fig. 6B**, **Table 1)**. For example, out of 64 ECM-related proteins, 62 genes in *S. A* infected-HCOs were down-regulated (e.g., COL3A1, COL1A1, MMP9, and MMP19) compared to non-infected ones (**Fig. 6B**, **Table 1)**. The infection also changed muscle contraction-related genes, such as upregulating MYL2 and CES1 (**Fig. 6C)**, and genes related to heart contraction, mainly including potassium ion channel protein-related genes (KCNE4 and KCNIP2) [34] and smooth muscle alpha-actin (ACTG2)[35], were all down-regulated. In *S. A* infection cases, the majority of ECM-related genes (> 90%; **Fig. 6B**) and muscle contraction-related ones (> 70%; **Fig. 6C**) were down-regulated compared to the healthy endocardium, indicating compromised tissue structure and motility by the bacterial infection.

Thus, we compared the data against the real-world transcriptomic data from human bacterial endocarditis (BE) studies (E-MEXP-1334)[36]. Notably, the differentially expressed (DE) genes of the COs overlapped with >566 genes (43.7%) of human DE genes (**Fig. S5D**) – which is even higher than the overlap between animal mouse and human models demonstrated by previous research[10]. Reactome analysis indicated that the overlapped genes included those related to ECM structure (**Fig. S5C, E**). Since ample clinical evidence suggests that bacteria-secreted toxins play key roles in mediating infective endocarditis[26a, 37], we investigated the proteomic change of *S. A* during their infection of HCOs, which revealed 701 and 145 significantly up-and down-regulated genes, respectively (**Fig. 6D**, **Table 2**). The expression correlation between *S. A* in planktonic status and those infecting HCOs was below 47%, while the inter-group correlation within *S. A* exceeded 97% and that within HCOs-SA exceeded 85% (**Fig. S5F**). Among the top 25 differentially secreted proteins (**Table 2**), exotoxins were the most significantly highly expressed proteins. The main exotoxins involved include leukocidin, fibrinogen-binding protein, and alpha-hemolysin (**Fig. 6E**). These toxins were clinically identified as the primary factor driving the pathological processes of *S. A* infection, leading to compromised heart contractions by destroying the cell and ECM structure[38],[39],[40]. The *S. A* could also penetrate inside the COs or DCM-COs after 48-h incubation (**Fig. 6F, G, H**)

Then, we inspected the structural changes of the COs after *S. A* infection (**Fig. 7A-D**). Indeed, the α-actinin distribution was significantly disrupted and became uneven after infection (**Fig. 7C** and **D**), as were MHC and troponin, though the overall integrity of the microtissue remained. These findings again underscored the significant impact of bacterial infection on the proteins related to tissue structure and myocardial contractility in the COs. Quantitative analysis of the images reveals that the structures of actin, calponin, and troponin (**Fig. 7E**) were severely disrupted after infection, while MHC (**Fig. 7F**) was relatively less affected.

**Figure 7.**
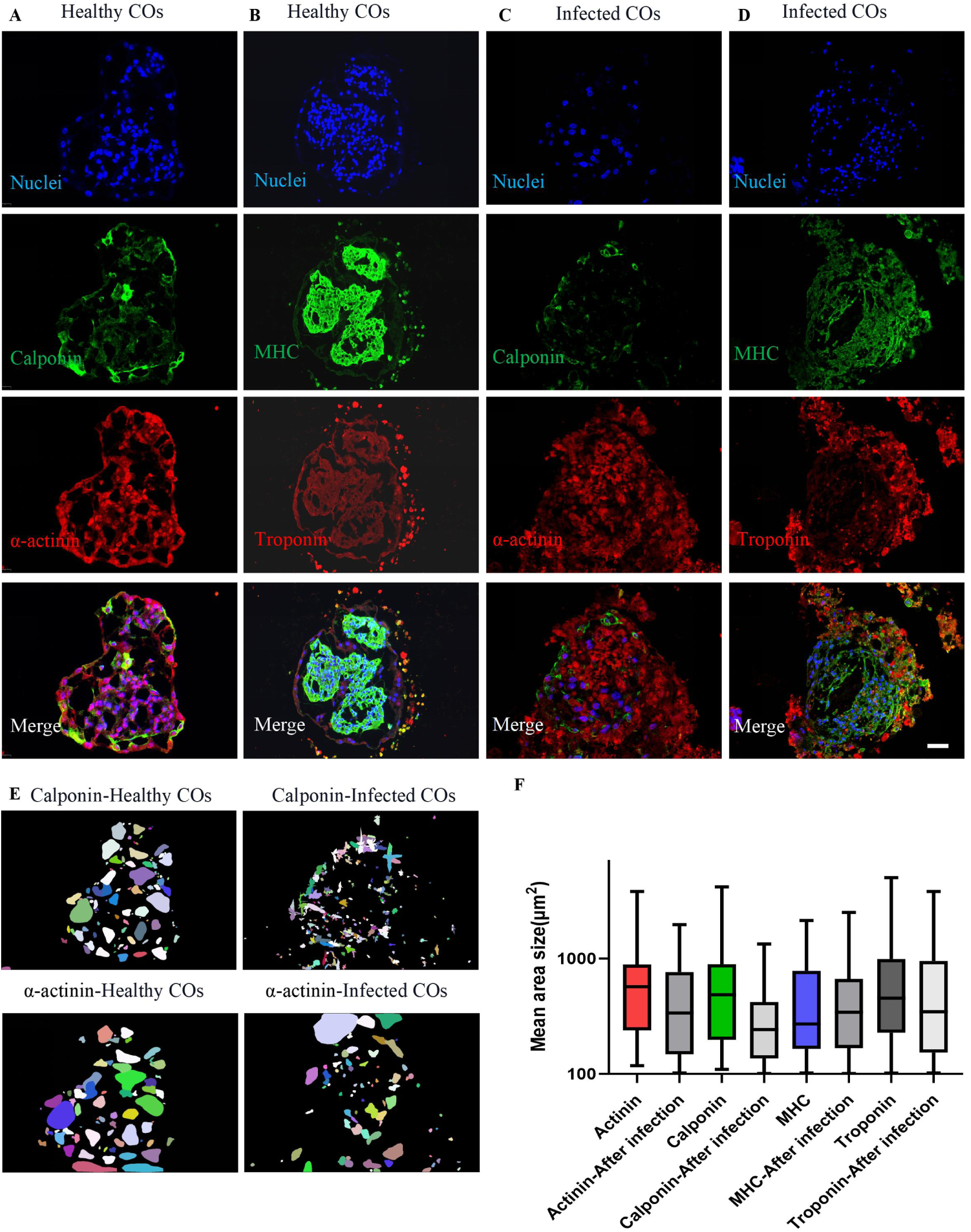
Bacterial infection reduced COs contraction by influencing myosin structure. (A) CLSM Observation HCO’s DAPI/ Calponin/ α-actinin localisation. The bar is 50μm. (B) CLSM Observation HCO’s DAPI/ MHC/ Troponin localisation. The bar is 50μm. (C) CLSM Observation HCO’s DAPI/ Calponin/ α-actinin localisation after infection after 24h co-culture with *S. A*. The bar is 50μm. (D) CLSM Observation HCO’s DAPI/ MHC/ Troponin localisation after infection after 24h co-culture with *S. A*. The bar is 50μm. (E) The distribution pattern of Calponin and α-actinin structures in HCOs before and after *S. A* infection, obtained through Stardist analysis. (F) Quantitative analysis of the formed areas of MHC, Troponin, Calponin, and α-actinin before and after infection to demonstrate the extent of structural disruption caused by the infection.

### Validation of increasing heart contractility as a potential strategy against bacterial infection

To validate that weakened contractility is key to facilitating bacterial infection of the myocardium, we established a mouse model with reduced cardiac function to observe *S. A* infection. We performed transverse aortic constriction (TAC) surgery to induce cardiac hypertrophy, as confirmed by electrocardiography with significantly lower heart rates (**Fig. 8A**). A total of 24 mice were divided into four groups (n=6 per group): i) mock group: mice in this group received a standard PBS and bacteria injection after TAC surgery, serving as a baseline for comparison; ii) amoxicillin group: mice in this group received the 50 mg/mL amoxicillin treatment and bacteria injection after TAC surgery; iii) enalapril group: mice in this group received the 10mg/mL Enalapril treatment and bacteria injection after TAC surgery; iv) sham group: mice in this group received PBS and bacteria injection without TAC surgery. The results of echocardiography also clearly validated this finding, with the sham group exhibiting a more normal electrocardiogram compared to the surgical group (**Fig. 8B**). The mice administrated with enalapril exhibited significantly higher cardiac output (Fractional shortening) than those in the mock group. At the same time, the amoxicillin treatment had a much weaker effect (**Fig. 8C**). Following surgery, all mice showed instances of tachycardia, which could be a compensatory response to the decreased cardiac function. After surgery and drug administration, we performed bacterial injections in each group. Subsequently, for three days, we administered the corresponding therapeutic drug once daily. The mice were euthanized three days later, and their hearts were collected to observe for any signs of bacterial infection (**Fig. 8D**).

**Figure 8.**
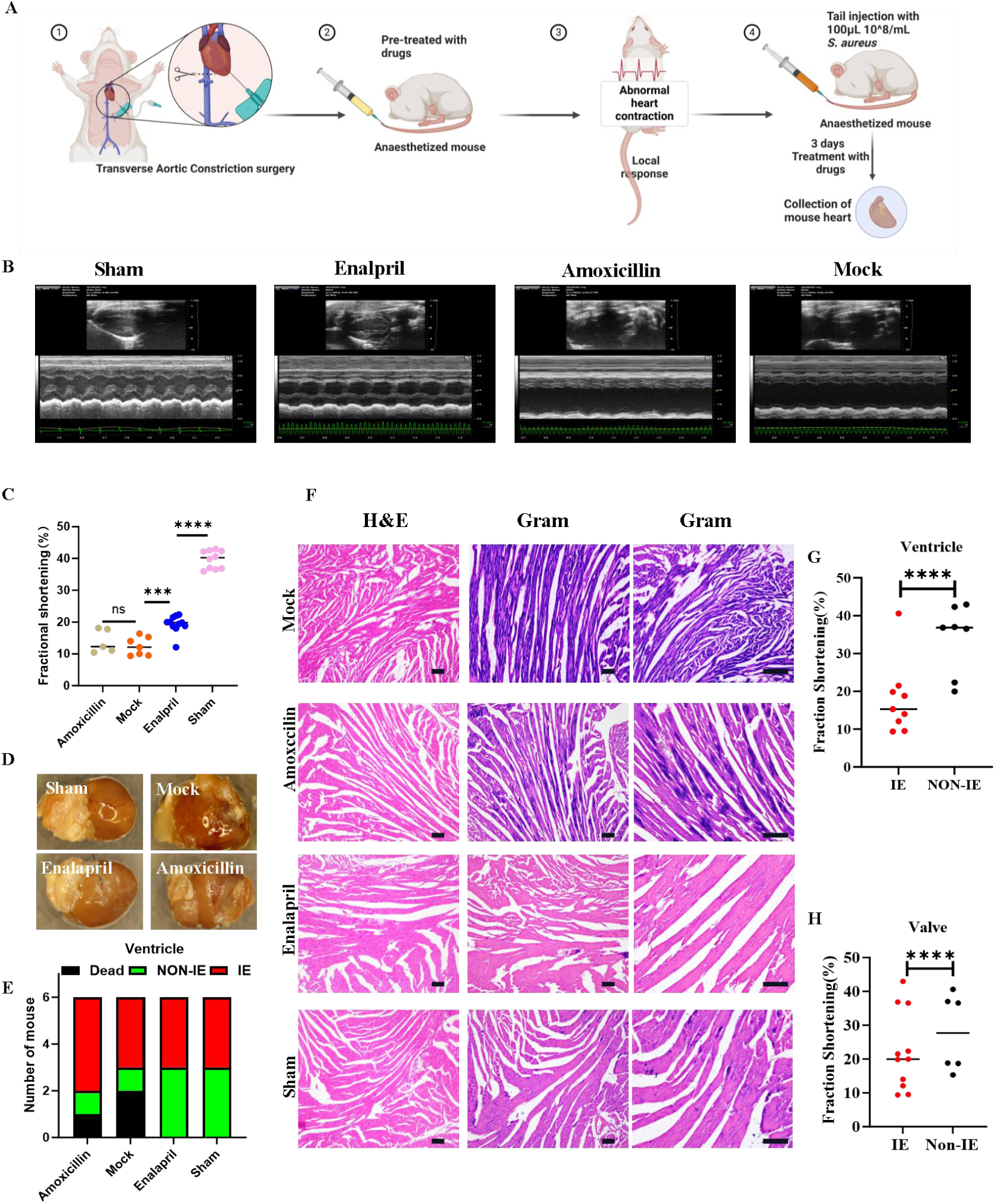
Dissecting the influence of contractility on *S. A* infection *in vivo* using mice model. (A) Experimental set-up: Mice were assigned into groups (Sham, Mock, Amoxicillin and Enalapril; n=6). All C57Bl/6 mice except the Sham group first received TAC surgery to induce reduced heart function. Different drugs were injected into mice for further experiments. (B) Electrocardiography of the mice from different experimental and Sham groups was measured two hours after drug administration. (C) Fraction shortening (%) of the mice from different experimental groups and Sham group measured by Electrocardiography at 2h after drug administration.\*\*\**P<*0.005 \*\*\*\**p*<0.001 (D) The heart specimens collected post-experimentation from different groups. (E) The number of deaths, IE on ventricle, and non-IE on ventricle among mice in different experimental groups. The infection status of decimals was confirmed through observation of cardiac bacteria vegetations and Gram staining under a microscope. (F) Aortic ventricle endocarditis in different experiment groups. Staining: haematoxylin and eosin (H&E) and Gram staining. The bar is 50 μm (G) Comparative analysis of the Fractal Dimension in mice with or without endocarditis in the cardiac ventricles in Enalapril, Mock and Sham groups. Significance:\*\*\*\**p*<0.001 (H) Comparative analysis of the Fractal Dimension in mice with or without endocarditis in the cardiac valves in Enalapril, Mock and Sham group. Significance:\*\*\*\**p*<0.001

Within three days following bacterial injection, one mouse from the amoxicillin group and the mock group died, while no deaths occurred in the enalapril group and the sham group (**Fig. 8E**). Enalapril effectively prevented bacterial infection on the ventricle while severe infections were observed in both the amoxicillin and the mock group (**Fig. 8F)**. Gram staining of the tissue specimen revealed that the ventricles of only one mouse in the Mock group and three in the sham group were infected. Amoxicillin treatment only spared one mouse (n=6) from infection, suggesting its ineffective prevention of *S. A* from ventricle colonization (**Fig. 8F**) However, the Enalapril treatment prevented three mice from ventricle infection (n=6), markedly reducing the probability of infection or death by 25% compared with the mock group and 16% compared to the amoxicillin group (**Fig. 8G**). The infection scenario in the valves was similar (**Fig. S6A**): enalapril significantly reduced the likelihood of valve infection or death by 16.6%, compared to the amoxicillin treatment and mock group (**Fig. S6B**). Notably, enalapril, aimed at increasing heart contractility but not inhibiting bacteria, performed better than the antibiotic amoxicillin in lowering the infection rate, reaching the same level as the sham group.

We performed statistical comparisons of FS (fractional shortening) and heart rate between mice infected with bacteria and uninfected mice, noting that the cardiac function of mice without ventricle infection was significantly higher by 25% than that in the infected mice (**Fig. 8G**). Also, the heart rate of mice increased significantly post-surgery, regardless of drug treatment, as a compensatory response to the decreased cardiac function. However, this elevated heart rate was not related to reduced bacterial infection (**Fig. S6C**). For valve infection; the FS and heart beat rate were both significantly increased (**Fig. 8H, Fig. S6D**) between mice without valve infection compared to those that were uninfected. These *in vivo* results validated our findings from engineering models correlating weakened myocardial contractility with increased bacterial infection, suggesting that increasing heart contractility, instead of heart beating rate, may become a new therapeutic approach for IE.

## DISCUSSION

Bacterial infection of the heart tissue can cause endocarditis with a high mortality rate, but the fundamental reason for the failure to resist microbial invasion of the myocardium remains unclear. Traditionally, it is understood that when events like open-heart surgery, trauma, or endothelial inflammation damage the endocardium integrity, bacteria manage to adhere to and invade the myocardium[41]. Several reports have also suggested that altered hemodynamics resulting from heart disease can influence cardiac bacterial infections[42]. However, one major underlying cause of altered hemodynamics[43], as well as a common underlying condition in many high-risk IE populations[44], heart failure, has been entirely overlooked.

In this study, by developing quantitative tools for analyzing bacterial-cardiac organoid interactions in a 3D, dynamic, clinically relevant setting, we discovered the significant role of cardiac contractility in preventing bacterial infection. Our created toolbox of live-cell image analysis integrated with the finite element analysis model, for the first time, unraveled the crosstalk between the self-propelled motion of the COs and the infection attempts by bacteria in different stages. The findings that patient-derived DCM-COs with weakened motility failed to resist bacterial adhesion – while healthy donor-derived HCO with strong contractile force managed to – suggest that bacterial infection of the heart is correlated with cardiac motility, even without pathological changes in cardiac valve structures. Interestingly, neither frequency nor pattern of the cardiac motion, as often speculated, played a significant role. Our findings provide fresh insights for possibly devising new contractility-enhancing therapies against endocarditis.

The heart is unique compared to other organs for its self-generated contractile force, which increases the difficulty of studying myocardial infection. The intrinsic contraction of the myocardial tissue and the pathogens’ motility should be considered interactively and quantitatively. Previous studies have provided valuable insights and tools for investigating similar and dynamic systems. For instance, Huebsch et al. developed mathematical models to assess the motion of COs36, and Mills et al. employed advanced systems to assess forces [45]. Several attempts were made to construct *in vitro* organ models to simulate infections [12, 46]. Sauvonnet et al.’s groundbreaking research explored the impact of intestinal peristalsis on bacterial infections, opening the door to studying the dynamics between organs and bacteria[47]. These studies have made significant contributions, but the relationship between cardiac cell motility and bacterial infection remained unexplored. We studied the impact of highly mobile cardiac organoids on bacterial infections by integrating physical, mathematical, and biological models. Our study has shed light on the significance of incorporating physical forces into the study of biological systems, with potential implications for understanding heart disease and related conditions.

We integrated cell tracking tools and FEM for live-cell image analysis, which showed unique advantages for analyzing COs’ contractility and COs-bacteria interactions in a real scenario. Utilizing image tracking, our established methods can readily obtain rich data about the motions without relying on any complex apparatus.

The FEM model we utilized here combines ease of operation with greater realism, as it takes into account the material properties of the cells themselves as well as the properties of the water field within the FEM system. The FEM system has been proven to be highly valuable in numerous cardiovascular studies[20b, 48]. We also conducted various statistical approaches to analyze the relationship between organoid movement patterns and adhesion status in different COs. The methodologies established in this study provide powerful tools for future research related to COs.

Importantly, our COs were derived from clinical samples with disease-relevant properties. We have innovatively found that COs with healthy cardiac muscle contractility were resistant to bacterial infection. DCM-COs, which had weak motility with a disease-related heartbeat pattern, did not exhibit this ability. After the contractility of COs was reduced by the toxins secreted during bacterial infection, COs could be infected by bacteria again. The functions of these toxins have been widely studied, and they play a very important role in infection, including adhesion, destruction of cell structures, and motility[37b, 37c, 49]. Our results agree with real clinical outcomes in either the contractile pattern or the contribution of toxins.

Several limitations of this study also inspire future directions. First, we only considered the self-propelled fluid motion of the organoids. Although the fluid motion in the body is also driven by the heart, the presence of the ventricles and blood vessels results in a different fluid form. Constructing a heart-like organ system using microfluidics can provide a more realistic fluidic environment for studying the interactions between bacteria and the host. Second, the role of immune cells is also crucial in understanding infectious diseases of any tissue. It is important to acknowledge the presence of resident macrophages, albeit in small numbers[50], among other immunocytes. At the same time, it is worth investigating how the heartbeat affects blood cells in the cardiovascular system during activation and migration. The different motility of the heart may guide the activation and adhesion of immune cells in a different direction). Third, the field still lacks research on bacteria and cells’ temporal and spatial gene expression during infection. To address this, more comprehensive techniques such as spatiotemporal transcriptomics and single-cell sequencing can provide detailed information about the gene expression of cells at different layers and the genomic expression of bacteria at different stages of infection. Finally, the degree of maturation of the organoids is also worth further optimization. In our study, we only observed the presence of some cavity-like structure within the organoids. This basic structure would be sufficient for simulating basic bacterial adhesion, similar to the organoids used by Richards et al[10]., in their simulation of myocardial infarction. For more accurately understanding the various stages of infection and the detailed host-microbe dynamics, more mature COs with well-developed lumen structures may also be necessary, as demonstrated previously by Hofbauer, P. et al[13a].

